# Transcriptomic plasticity in hybrid schistosomes can contribute to their zoonotic potential

**DOI:** 10.1101/2024.12.16.628740

**Authors:** Nelia Luviano Aparicio, Eglantine Mathieu-Bégné, Julien Kincaid-Smith, Olivier Rey, Marion Picard, Cristian Chaparro, Jean-François Allienne, Anne Rognon, Bruno Polack, Isabelle Vallée, Myriam Thomas, Jean-Jacques Fontaine, Jérôme Boissier, Eve Toulza

## Abstract

Hybrids between *Schistosoma haematobium* and *S. bovis* are linked to both human and animal infections, highlighting the complex interspecies interactions that contribute to the spread of schistosomiasis. Additionally, *S. bovis* can infect multiple ruminant hosts, facilitating cross-species transmission and increasing the risk of zoonotic outbreaks. In this study, we investigated transcriptomic plasticity as a potential mechanism enabling hybrid schistosomes to adapt to alternative definitive hosts. We focused on two contexts: 1) introgressed *S*.

*haematobium × S. bovis* hybrids, which demonstrated higher virulence in sheep compared to parental *S. bovis*, and 2) *S. bovis* infecting different host species. Our analysis uncovered 366 differentially expressed genes (DEGs), representing 4% of the total protein-coding genes, between introgressed hybrids and parental *S. bovis* in sheep. We also identified transcriptomic changes in *S. bovis* across different mammalian hosts (hamster and sheep), with around 30% of the total genes differentially expressed, demonstrating that *S. bovis* parasites display a high transcriptomic plasticity, allowing them to infect different definitive hosts. Shared enriched biological processes during introgression and host change include nuclear-transcribed mRNA catabolic processes, inner mitochondrial membrane organization, microtubule-based movement, response to endoplasmic reticulum stress, and sensory perception. These findings suggest that transcriptomic plasticity in *S. bovis* and hybrid worms enhance their ability to adapt and infect diverse host species, potentially increasing their zoonotic potential. This raises concerns for schistosomiasis epidemiology, as this plasticity could expand the parasite’s transmission capacity and complicate control efforts.

## Introduction

Hybridization can significantly influence the ecology and evolution of host-parasite interactions. The viable offspring resulting from hybrids between two species might also backcross with one of their parental species. If these backcrossed offspring continue to reproduce with the same parental species, this can result over time in the lasting transfer of DNA from one of the species into the other, a process known as introgressive hybridization or introgression (Aguillon et al. 2022). Introgression can promote gene flow between two species, increasing the genetic variability and fitness of one or the other species and facilitating adaptation to new environments. Both host and parasite introgressive hybridization have been shown to impact parasite virulence, transmission, and host specificity. However, research on hybridization has primarily focused on how hybrid hosts are affected by parasites, treating parasites as a selective force in host evolution. This underscores the importance of considering that parasites also undergo their own evolutionary dynamics (Detwiler and Criscione 2010).

In this work, we focus on schistosome parasites, which serve as an excellent model for studying introgression. Schistosomes are parasitic blood flukes that develop and grow in the vascular system of mammal definitive hosts. They are responsible for causing the bilharzia or schistosomiasis, a neglected tropical disease (NTD) and the second most important parasitic disease after malaria. There are over 230 million people requiring treatment, of which the majority live in Africa (Mujumbusi et al. 2023). Schistosomiasis is a waterborne disease. Schistosome parasites first infect at the miracidiae stage an intermediate mollusk host in which the parasite asexually multiplies. Cercariae are then released and can infect the mammal definitive host when this later has contact with water. Sexual reproduction occurs in the definitive host and, depending on the schistosome species, eggs are released through urine or feces in water so as the cycle can continue.

While numerous schistosome species exist, their barriers are often maintained through differences in ecology, host specificity, and evolutionary history (J. Kincaid-Smith et al. 2021). However, hybridization between closely related species can happen when parasites infect the same host (Webster et al. 2013). For instance, hybrids between *Schistosoma haematobium* and *S. bovis* have been detected in humans across several African countries, including Benin, Ivory Coast, Mali, Niger, Senegal (Leger and Webster 2017; Angora et al. 2020; Onyekwere et al. 2022). These hybrids have also been found in cattle in Benin which evidences that these hybrids are present in livestock of sub-Saharan Africa (Savassi et al. 2020). Furthermore, *Schistosoma bovis*, as well as hybrids between *S. bovis* and *S. curassoni*, have been observed in sheep in the field (Rollinson et al. 1990). By conducting laboratory infections of hybrids between *S. bovis* and *S. haematobium* in sheep, we can gain a better understanding of the potential for hybridization in this host and its phenotypic and epidemiologic implications (Stothard et al. 2020).

*Schistosoma bovis* infects numerous ruminant species including cows and sheep (Makundi et al. 1998; Ferreras et al. 2000) and can also use some rodents as definitive hosts including the hamster (*Mesocricetus auratus*) which is commonly used as definitive hosts to maintain the parasite under laboratory conditions (Southgate et al. 1985). Previous studies have examined gene expression in *S. bovis × S. haematobium* hybrids, but these have been limited to infections in hamsters (Mathieu-Bégné et al. 2024). It remains unclear whether similar patterns occur in natural host or what changes in the parasite’s transcriptomic machinery might facilitate long-term host adaptation in backcrossed individuals. This has significant implications given that most experimental research on schistosomes relies on host species rarely encountered in the field.

Both *S. haematobium* and *S. bovis* are phylogenetically close and share freshwater snails of the genus *Bulinus* as their intermediate host (Teukeng et al. 2022). The study of the molecular variation of *S. haematobium* among Africa brought to light genomic signatures that correspond to species other than *S. haematobium,* suggesting the occurrence of introgression events (Stroehlein et al. 2022). Indeed, it was demonstrated that most *S. haematobium* African populations show signatures of genomic introgression from *S. bovis* (Rey et al. 2021) .

In the case of *S. haematobium × S. bovis* hybrids, introgressive hybridization may enable the rapid acquisition of new traits, potentially facilitating adaptation to novel hosts and environments. RNA expression analysis of these hybrids has shown overexpression of processes linked to heterosis, particularly in females, which may be associated with increased reproductive potential (Mathieu-Bégné et al. 2024).

The zoonotic potential of these introgressed hybrids warrants attention in epidemiological fieldwork, including diagnostics and treatment strategies, as they may display enhanced virulence and a broader host range compared to their parental species (Julien Kincaid-Smith et al. 2021). This was confirmed by a recent study which found that introgressed *S. haematobium* × *S. bovis* worms can infect sheep, whereas pure *S. haematobium* parasites cannot (Polack et al. 2024). This suggests that schistosomes possess a molecular mechanism that allows host change, particularly when hybridized with another species (Mason, 2016), raising public health concerns about their ability to exploit new hosts and regions (Mathieu-Bégné et al., 2022; Thompson et al., 2009). Despite this, there is limited knowledge about the molecular machinery allowing parasites to successfully infect alternative definitive host species, and this knowledge exists mostly for parasitic insects (Hardy and Otto 2014; Mason 2016) but is absent in other lineages such as Schistosoma In this study, we analyzed the transcriptomes of introgressed *S. haematobium × S. bovis* worms, which displayed increased virulence in sheep (Polack et al., 2024) and investigated gene expression changes associated with host change in *S. bovis*. The goal was to uncover biological processes that might enhance the zoonotic potential of both *S. haematobium × S. bovis* hybrids and *S. bovis*. We hypothesize that: 1) introgressed *S. haematobium × S. bovis* parasites exhibiting higher virulence in sheep compared to parental *S. bovis* undergo significant transcriptomic changes; 2) host change in *S. bovis* is accompanied by substantial transcriptomic plasticity that facilitates adaptation to different host species and 3) the transcriptomic changes associated with both introgression and host change involve common biological pathways

## Material and Methods Ethics approval

Sheep infections with *S. bovis*, *S. haematobium* and hybrid cercarias were conducted in the Veterinary School Ecole Nationale Vétérinaire d’Alfort in Maisons-Alfort, France. The animal facility possesses agreement E940462 delivered by the French Ministry of Higher Education, Research and Innovation. The sheep experiment was registered by the French Ministry of Higher Education, Research and Innovation, and approved by an independent ethics committee (APAFIS#23786-2020012416159143 v2) Worms were collected and frozen in liquid nitrogen and stored at -196°C for subsequent analysis.

## Parasite and host strains

The *S. haematobium* strain originated from infected patients (with ethical clearance) in the Southeast part of Cameroon (Barombi Kotto lake) isolated in 2015 and maintained in the IHPE laboratory using *Bulinus truncatus* and the golden hamster *(M. auratus*) as intermediate and definitive hosts. The *S. bovis* strain originated from the Spanish laboratory of parasitology of the Institute of Natural Resources and Agrobiology in Salamanca and was maintained in the IHPE laboratory using *B. truncatus* and *Planorbarius metidjensis* as intermediate hosts and the golden hamster *M. auratus* as definitive host.

## Experimental infection of the targeted definitive hosts

Hamster infections with the different parasite strains including the pure *S. bovis strain,* the F1 and F1’ hybrids and the backcross to *S. bovis* and *S. haematobium* were conducted as previously described in Mathieu-Bégné et al., 2024. The experimental infections in sheep with the different parasite strains were conducted at the Maisons-Alfort laboratory as previously described in (Polack et al. 2024). Briefly, cercariae from the different laboratory parasite strains were used to infect 3 series of sheep for each parasite strain: parental *S. bovis* and *S. haematobium* strains, F1 (and F1’) parasites, and backcrossed parasites (Table 1).

**Table 1.** Crosses and backcrosses between *S. bovis* and *S. haematobium* used in this work (only adult parents are represented) using *B. truncatus* and the golden hamster or sheep as intermediate and definitive host respectively.

| Male Parent | Female Parent | Cross type |
| --- | --- | --- |
| *Schistosoma bovis* | *Schistosoma bovis* | Pure *S. bovis* |
| *Schistosoma haematobium* | *Schistosoma haematobium* | Pure *S. haematobium* |
| *Schistosoma bovis* | *Schistosoma haematobium* | F1 Hybrid (*S. haematobium* × *S. bovis*) |
| *Schistosoma haematobium* | *Schistosoma bovis* | F1 ‘Hybrid (*S. bovis* × *S. haematobium*) |
| *Schistosoma haematobium* | F1 Hybrid | Backcross *S. haematobium* |
| *Schistosoma bovis* | F1’ Hybrid | Backcross *S. bovis* |

Hamsters and sheep were infected following a natural percutaneous immersion method. Three control sheep were exposed to the pure strains: *S. bovis* and *S. haematobium*. The same number of cercariae (1500) was used to infect each sheep and 300 cercariae for hamsters. For each parasite line, a minimum of 12 snails exposed to 5 miracidia produced the cercariae for sheep and hamsters’ infection. Sheep infection was done under anesthesia (ketamine and diazepam) at the level of a forearm previously wet with warm water. Sheep euthanasia was performed by deep anesthesia with a combination of xylazine and ketamine, followed by bleeding the animals to collect blood. Sheep autopsies were conducted under the supervision of a European College of Veterinary Pathologists board member. All viscera were examined, and the vessels of the small intestine, cecum, colon, rectum, and bladder were examined by candling with a strong light.

Adult male and female schistosomes were recovered from experimental infections of sheep and hamsters for each cross type. The goal was to collect adult parasites resulting from experimental infections as a primary resource for subsequent transcriptomic analyses. In sheep, the worms were isolated by opening the vein with a scalpel and carefully extracting the live schistosomes using forceps. In hamsters, worms were recovered through portal vein perfusion. Male and female worms were carefully separated to avoid cross-sex contamination and pooled into groups of 12 individuals per condition and sex. Three biological replicates were prepared for each group, resulting in a total of 48 samples.

## RNA extraction and transcriptome sequencing procedure

Total RNA extractions were performed using the Qiagen RNeasy mini kit. Briefly, pools of 12 frozen adult worms in 2ml microtubes were grounded with glass beads (Sigma, reference G4649) using a blue pestle RNase free in the case of females, and regarding males, these were grounded with a different method (due to the higher size) using zirconium beads and the Retsch MM400 cryobrush (2 pulses at 30Hz for 15s). After extraction following manufacturer’s protocol, total RNA was eluted in 44 μl of RNase-free water. DNase treatment was then performed using Thermofisher Scientific Turbo DNA-free kit.

Quality and concentration were assessed by spectrophotometry with the Agilent 2100 Bioanalyzer system and using the Agilent RNA 6000 nano kit. cDNA library construction and sequencing were performed by the Bioenvironment sequencing platform (University of Perpignan, France) with the NEBNext Ultra II directional RNA library preparation kit following the manufacturer’s protocol on 300 ng of total RNA per sample. Sequencing was performed in 75 bp single end on an Illumina NextSeq550 instrument at the Bio-Environment platform (UPVD).

### Transcriptome analyses

The RNA sequencing data from *S. bovis* infecting hamster that was used to analyze transcriptomic adjustments associated with host change were recovered from the Bioproject PRJNA491632, for which we used six samples from the homospecific pairing, 3 males and 3 females (Biosamples: SAMN10081090, SAMN10081089, SAMN10081088, SAMN10081087, SAMN10081086 and SAMN10081085).

Bioinformatic analyses were performed on the Galaxy instance of the IHPE laboratory. Raw reads were subjected to quality assessment using FastQC tool version 0.72 (Andrews 2010). We trimmed reads based on quality (phred quality score threshold<20) using trim-galore tool version 0.6.3 (Krueger 2012). Processed reads were mapped to the *S. haematobium* reference genome version 3 since it has a better genomic resolution than the *S. bovis* genome (Stroehlein et al. 2022) using RNA STAR mapper version 2.7.8 (Dobin et al. 2013).

Counting of reads per transcript was done using htseq count tool (Version 0.9.1) and using the gene transfer file from *S. haematobium* as reference transcriptome (schistosoma_haematobium.PRJNA78265.WBPS18.annotations.gff3 downloaded from Worm Base Parasite).

Differential expression analyses were done with the R package DESeq2 (Love et al. 2014), low counts were filtered (<10), and genes differentially expressed were identified by setting a *adjusted p-value* < 0.05 and log_2_FC > 1. We performed Principal Components Analysis for all the samples to identify clusters between samples. To identify gene expression modifications associated with hybridization, we compared the gene expression of parental *S. bovis* to those of the first-generation hybrids (F1 and F1’). To identify gene expression modifications associated with introgression, we compared the gene expression of parental *S. bovis* to those of the backcrosses male F1 x female *S. bovis* that displayed maximal infection rates in sheep (Pollack et al. 2024). Since the differences between sexes were stronger than those between genetic introgression levels (see results), we conducted these analyses for each sex separately. To specifically address which modifications in gene expression are associated with host change, we compared the transcriptome between the same *S. bovis* strain infecting the hamster and the sheep and accounting for a possible sex effect. To this aim, we compared three pools of 12 males, and three females collected from each of the two definitive hosts (i.e. hamster and sheep). One female *S. bovis* collected from sheep was discarded from the analysis as it was identified as an outlier. For all pairwise comparisons, we specified which samples to compare using the ’contrast’ parameter in the DESeq2 software. This allowed us to analyze the differences in gene expression between the selected samples. It is worth to mention that we could not analyze *S. haematobium* worms because they were not capable of infecting sheep.

The reference transcriptome assembly was converted from GTF to FASTA format using gffread (version 2.2.1.3), with the reference genome provided as input (Trapnell et al. 2010).

This reference transcriptome in FASTA format was used as input in Orson, a nextflow based workflow for transcriptome functional annotation developed by the Bioinformatics Core Facility of Ifremer, the French National Institute for Ocean Science and configured on DATARMOR supercomputer (https://gitlab.ifremer.fr/bioinfo/workflows/orson). Merged XML file output from Orson workflow was then used as input in OmicsBox bioinformatics software v 1.3.11 (OmicsBox, 2019) to execute GO mapping and GO annotation.

GO enrichment analysis of differentially expressed genes was performed using the R package RGBOA, available on GitHub (GO-MWU, https://github.com/z0on/GO_MWU), we used as input a csv file containing all genes with binary values (1 when the gene was significantly over or under-expressed and otherwise 0) and we used the Fisher’s exact test mode, we selected enriched biological processes with a False Discovery Rate FDR<0.05 as used in another study (Wright et al. 2015). When the fisher test did not reveal significant enrichment of GO terms, we selected the top 10 of genes over and under-expressed for functional investigation. The software Revigo was used to remove redundant GO terms from the long lists obtained in the GO enrichment analysis and to illustrate representative enriched GO terms in a dot-plot (Supek et al. 2011).

## Results

### Transcriptomic analysis in hybrids and backcrossed schistosomes compared to parental S. bovis

On average we obtained 27,570,576 raw reads per sample after sequencing, and 27,283,136 reads per sample after quality trimming. On average 83% ±3.5 of the reads mapped to the reference genome of *S. haematobium* (V3) which consists in 9431 protein-coding genes, the percentage of mapping per sample is provided in the Supplementary file 1, Table S1. No significant differences in mapping % was observed depending on the species/hybrid line (p = 0.5, Kruskal-Wallis test), highlighting the high similarity between the two genomes, at least in the coding regions. Based on their overall gene expression patterns, parasites first clustered according to their sex along the first PCA axis that represented 48% of the overall transcriptomic variance among individuals and then according to their introgression level along the second PCA axis (representing 14% of the variance) (Figure 1A).

**Figure 1.**
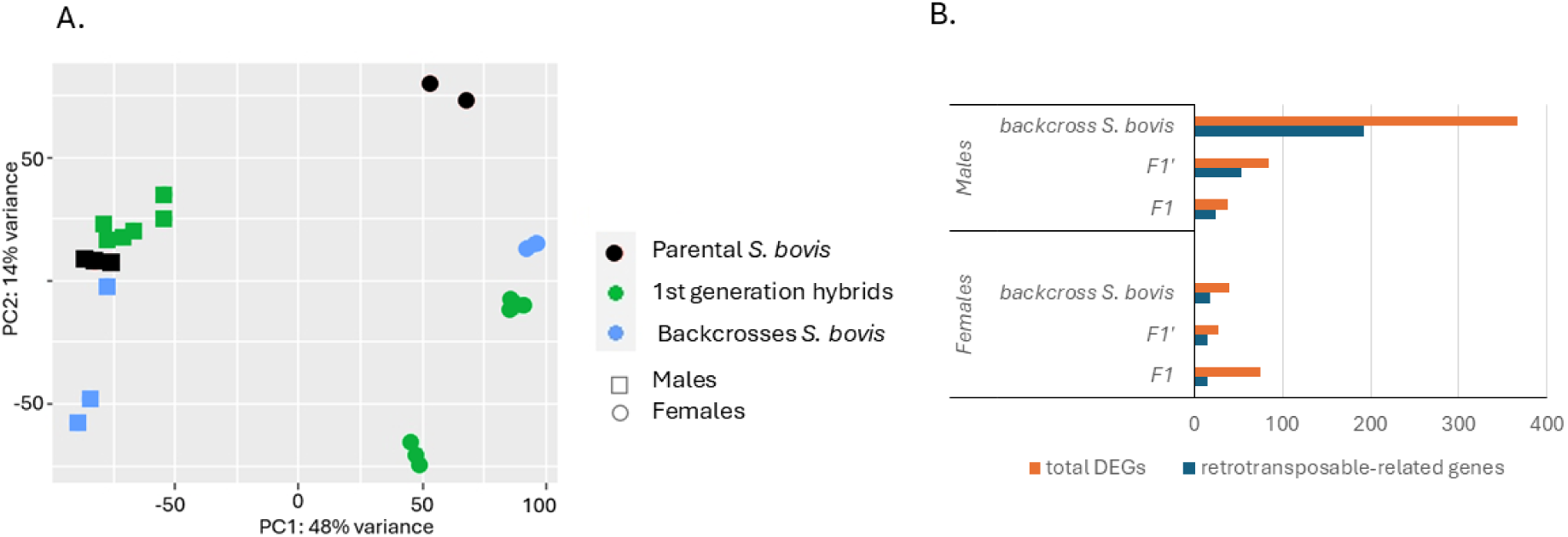
A. Principal component analysis (PCA) plot showing the distribution of transcriptomes of schistosomes of the two sexes with different levels of genetic introgression. Each dot in the plot represents a sample, and the different colors represent the different levels of genetic introgression The form indicates the sex, with squares representing males and circles representing females. **B.** Number of genes differentially expressed related to retrotransposable elements across all comparisons to *S. bovis*.

As sex was the first driver of transcriptomic differences, we next analyzed male and females separately. In females, we found 76 differentially expressed genes (DEGs) between *S. bovis* and first-generation hybrids from F1 crossing (♀*S. haematobium* × ♂ S. *bovis*), 35 over-expressed and 41 under-expressed in *S. bovis* compared to F1. For the comparison between *S. bovis* and F1’ (♀ S. *bovis* ×♂ *S. haematobium*), we found 28 DEGs, 21 over-expressed and 7 under-expressed (Supplementary file 2, Table S1-S2). By comparing females of *S. bovis* and those of the backcross with *S. bovis*, we found 40 DEGs, 20 of which were over-expressed and 20 under-expressed in the backcrossed compared to pure *S. bovis* females.

Concerning the comparison between males, we found 38 DEGs between *S. bovis* and F1, 18 over-expressed and 20 under-expressed. In the comparison between males *S. bovis* and F1’, we found 85 DEGs, 68 over-expressed and 17 under-expressed (Supplementary file 2, Table S3- S4). In the males, comparison between S*. bovis* and backcross *S. bovis* we found 366 DEGs, 260 were over-expressed and 106 under-expressed in backcrossed compared to pure *S. bovis* males (Supplementary file 2, Table S5-S6).

Over all the transcripts identified in these DEG analyses, we found many transcripts related to domains of retrotransposons. This represented more than 60% of the DEGs in first generation hybrid (F1 and F1’) males compared to pure *S. bovis* males (Figure 1B). Similarly, more than 50% of DEGs were related to retrotransposable elements in males from F1 strains backcrossed with *S. bovis* (Figure 1B). Despite fewer DEGs observed between female parasites from the three strains differing in introgression levels (i.e. pure *S. bovis*, F1/F1’ and backcross) similar percentage of retrotransposable elements were observed among the overall DEGs (i.e. 45% in backcrossed females, 54% in F1’ females) although less in the F1 females (20%) (Figure 1B).

The complete annotations of all the differential expressed genes are provided in the Supplementary file 2, Table S7.

### Transcriptional plasticity of the parasite *S. bovis* infecting two different definitive hosts

We explored the transcriptional plasticity associated with host change in the parasite *S. bovis* and the biological processes in the responsive genes are involved. On average 80% ±7.4 of the reads mapped to the reference genome of *S. haematobium* (V3), the percentage of mapping per sample is provided in the Supplementary file 1, Table S1.

The Principal Component Analysis showed that based on their whole gene expression patterns, parasites clearly clustered according to their definitive hosts (i.e. hamster versus sheep) along the first axis and according to their sex along the second axis; these two axes capturing 72% (respectively 52% and 25%) of the overall variability between samples transcriptomes (Figure 2).

**Figure 2.**
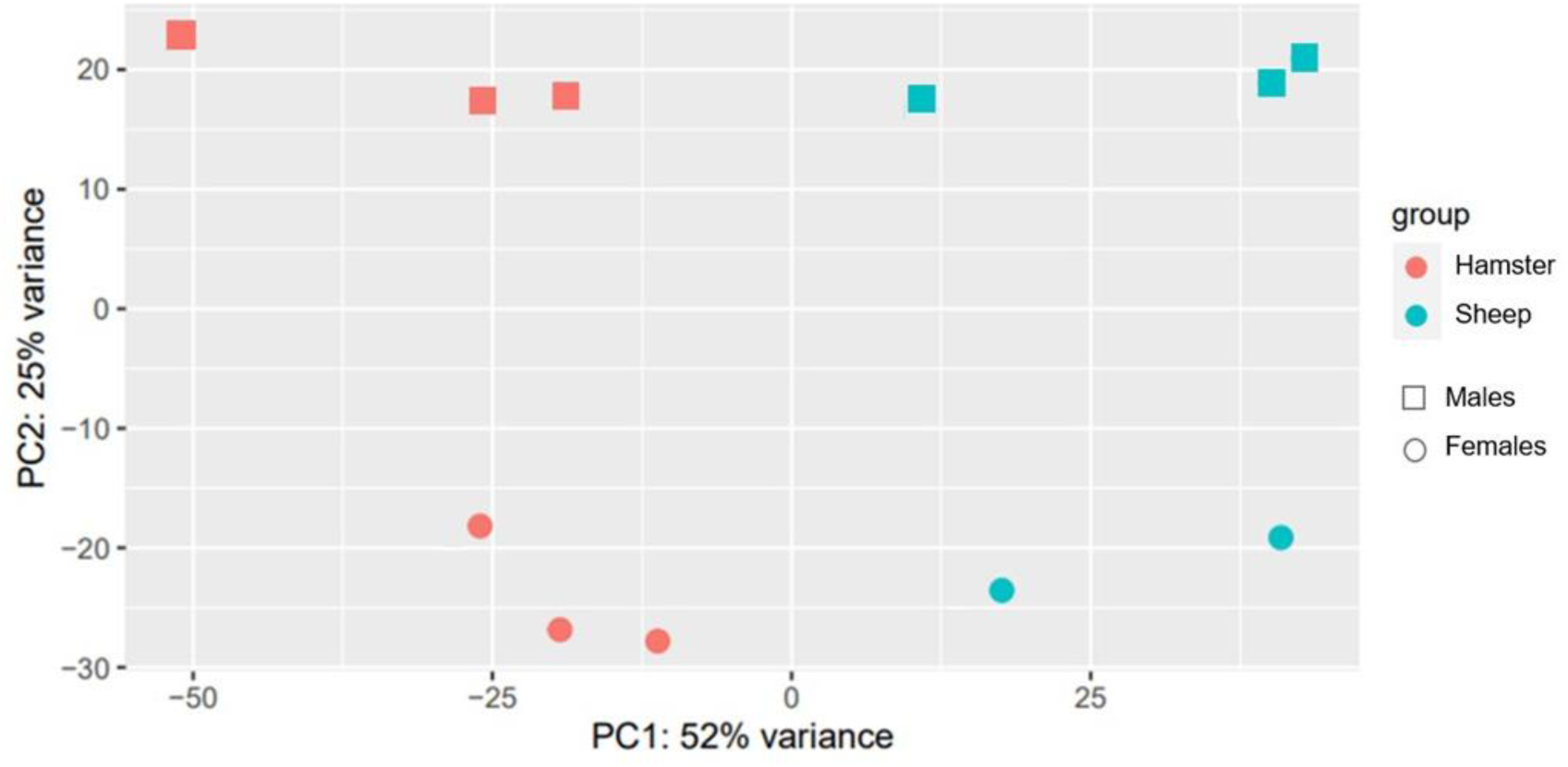
PCA analysis plot of male (square) and female (circle) *S. bovis* worms collected from sheep (blue) and hamsters (red).

In females, we found 2488 DEGs between parasites infecting sheep *vs.* hamster hosts, which represented 26% of the total number of genes, 1070 were over-expressed and 1418 under-expressed. In males, we found 2891 DEGs between parasites infecting sheep *vs.* hamster hosts, which represent 30 % of the total number of genes, 1192 being over-expressed in sheep and 1699 being under-expressed.

### GO enrichment analysis of introgressed worms and *S. bovis* from different hosts

#### Backcross S. bovis vs. parental S. bovis

To investigate the biological processes associated with gene expression differences during introgression, we performed a GO enrichment analysis in the differential expressed genes between males *S. bovis* and introgressed backcross worms (♂F1 *× ♀ S. bovis*). These analyses focused on comparisons between groups exhibiting the most divergent infection phenotypes, with significant processes identified based on an FDR-adjusted p-value < 0.05.

In males from the backcross (F1 *× S. bovis*), over-expressed genes were significantly enriched for processes related to nuclear-transcribed mRNA catabolic process, inner mitochondrial membrane organization, and response to endoplasmic reticulum stress. Conversely, under-expressed genes were enriched for processes including sensory perception, fucosylation, cilium/flagellum-dependent cell motility, neuropeptide signaling pathway, non-motile cilium assembly, microtubule-based movement, oligosaccharide biosynthetic process, and glucosylceramide metabolic process (Figure 3, first panel).

**Figure 3.**
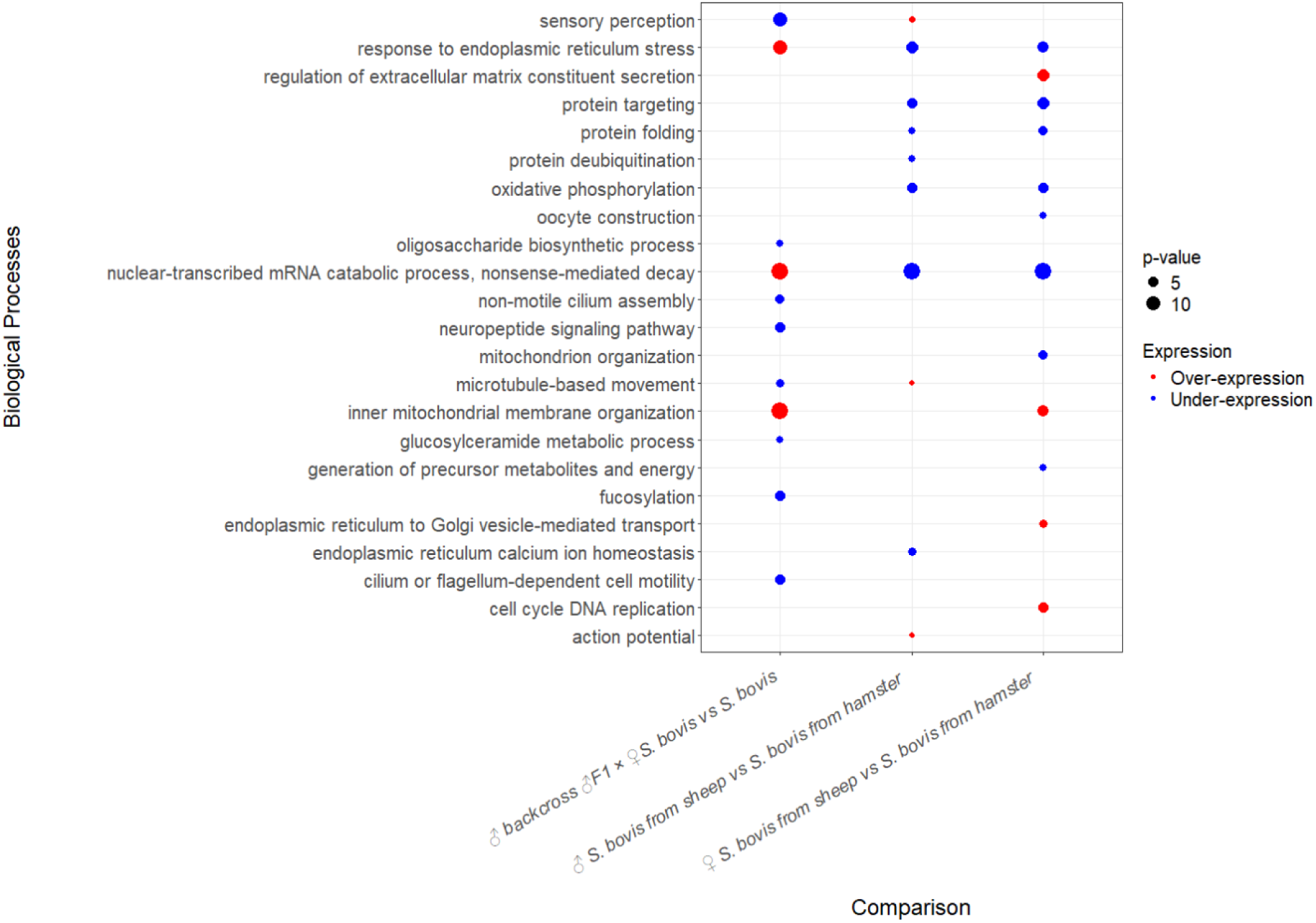
Dot plot illustrating the representative biological processes (y-axis) across experimental comparisons (x-axis). Dot size indicates the transformed significance level (p-value converted to percentages for improved visualization), with smaller percentages denoting greater significance. Dot color represents differential gene expression levels, where red indicates over-expression and blue indicates under-expression. The x-axis reflects specific experimental comparisons, highlighting the biological processes shared across conditions.

#### S. bovis infecting sheep vs. S. bovis infectin hamster

In males of *S. bovis* from sheep, over-expressed genes were enriched for processes such as action potential, sensory perception and microtubule-based movement. Under-expressed genes in this comparison were significantly associated with nuclear-transcribed mRNA catabolic process, nonsense-mediated decay, response to endoplasmic reticulum stress, protein targeting, protein folding, protein deubiquitination, oxidative phosphorylation, and endoplasmic reticulum calcium ion homeostasis (Figure 3, middle panel).

In females of *S. bovis* infecting sheep, over-expressed genes were linked to regulation of extracellular matrix constituent secretion, cell cycle DNA replication, inner mitochondrial membrane organization, and endoplasmic reticulum to Golgi vesicle-mediated transport. In contrast, under-expressed genes in females infecting hamsters were enriched for nuclear-transcribed mRNA catabolic process, nonsense-mediated decay, mitochondrion organization, protein targeting, protein folding, response to endoplasmic reticulum stress, oxidative phosphorylation, oocyte construction, and generation of precursor metabolites and energy (Figure 3, third panel).

## Discussion

In this study, we investigated the differential gene expression in schistosome hybrids (*S. haematobium × S. bovis*) and backcrossed worms compared to parental *S. bovis*. We also examined the transcriptomic changes in *S. bovis* infecting sheep compared to those infecting hamsters. Our findings offer valuable insights into the transcriptomic plasticity that may facilitate the ability of *S. bovis* and *S. haematobium × S. bovis* hybrids to exploit diverse host environments. Collectively, these findings underscore the role of transcriptomic plasticity in shaping the evolutionary trajectories of schistosome hybrids. This plasticity allows the parasites to navigate physiological and immunological challenges, expand their host range, and increase their zoonotic potential

## Differential expression in first generation hybrids

In first-generation hybrids less than 1% of the genes are differentially expressed compared to pure *S. bovis* parasites. These results are consistent with our previous results indicating that the most prevalent profile of gene expression in hybrids is at intermediate level between expression of each parental strain (Mathieu-Bégné et al. 2024). This highlights the high genomic compatibility between *S. bovis* and *S. haematobium*, which arises from their shared evolutionary history and limited divergence.

Differential gene expression analyses between first-generation hybrids and parental *S. bovis* revealed that differentially expressed genes (DEGs) were associated to retrotransposable elements. Retrotransposable elements were also found to be differentially expressed in hybrids of *S. bovis* and *S. haematobium* infecting hamsters, suggesting that these elements are mobilized independently of the host species (Mathieu-Bégné et al. 2024). Even if the transposable elements are not annotated neither in *S. bovis* no in *S. haematobium*, we found homologous sequences to some retrotransposable elements that have been previously identified in transcriptomic analysis of schistosome worms (Jolly et al. 2007; J. Kincaid-Smith et al. 2021).

The percentage of differentially expressed genes linked to transposon activity was found to vary between 20% in the comparison of female *S. bovis* and F1 hybrids, but it went up to 70% when comparing male *S. bovis* and F1 hybrids. Moreover, we also found that shared DEGs in males were all related to retrotransposon elements.

Our results support the hypothesis that hybrid offspring experienced a genomic shock due to the combination of two genome species, which results in genomic rearrangements such as transposition and chromosomal reorganization (McClintock 1984; Dion-Côté et al. 2014; Drouin et al. 2021; Mathieu-Bégné et al. 2024). These retrotransposons have the potential to influence the genetic identity and genome size. They can also impact regulatory networks, gene expression, and lead to genome rearrangements, ultimately providing important source of phenotypic and genetic diversity available for ecological and evolutionary processes (Rey et al. 2016; Serrato-Capuchina and Matute 2018).

In conclusion, these results suggest that even rare hybridization events—such as those likely occurring between *S. bovis* and *S. haematobium* (Platt et al. 2024)—can lead to significant genomic innovations. These events can facilitate the introgression of specific genomic segments from one species into another and mobilize transposable elements within this new lineage.

## Differential expression found in introgression

We found 366 DEGs, representing 4% of the total coding genes (9431) when comparing the transcriptome of the hybrid strain backcrossed with *S. bovis* (males F1× females *S. bovis*) that showed an increased virulence on sheep to the transcriptome of parental *S. bovis*. This is a higher number of DEGs than those found in first-generation hybrids compared to the *S. bovis* parental strain. Thus, even though backcrossed parasites display a genetic background that is getting closer to that of the parental *S. bovis* strain compared to F1 hybrids, their transcriptome continues to derive from that of the parental strain. This sustained transcriptomic differentiation could be a key factor driving the increased virulence observed in these hybrids. It implies that even subtle genetic introgression may have significant functional consequences on parasite biology, potentially affecting traits like host specificity and immune evasion, thus contributing to their enhanced virulence. It has already been reported that introgressed individuals exhibit increased virulence (Polack et al. 2024). In this work, we focused on elucidating the molecular processes that underly this enhancement in virulence. Understanding these mechanisms is crucial, as it may provide insights into how genetic interactions influence pathogenicity and inform future research on the evolution of virulence in these organisms.

In the over-expressed genes in the backcrosses versus the parental strains a prominent enrichment for processes related to nuclear-transcribed mRNA catabolic process and inner mitochondrial membrane organization was observed. The enrichment of mRNA catabolic process indicates a potential increase in RNA surveillance and turnover. This could reflect regulatory mechanisms for fine-tuning gene expression in response to genetic introgression (Buccitelli and Selbach 2020). The enrichment of mitochondrial organization suggests an increased metabolic activity in introgressed worms (Giacomello et al. 2020).

The biological processes enriched among the genes under-expressed in males from backcross F1 × *S. bovis* compared to males of the *S. bovis* parental strain were sensory perception and neuropeptide signaling pathway. This suggests that schistosomes from the backcrossed strain, might have altered sensory perception and neuronal signaling, potentially impacting their interactions with the host. Notably, processes associated with ciliary function and flagellum-dependent cell motility, are also enriched. Cilia are essential organelles that play a critical role in sensing environmental cues and facilitating movement (Nachury 2014). The underexpression of genes involved in these processes could have implications for the parasite’s ability to navigate and interact with the host vascular environment.

Schistosomes adult worms have a variety of sensory structures that allow them to respond to changes in levels of chemicals and nutrients in the host vascular system. These include various chemoreceptors and membrane channels. We found G protein-coupled receptors that have previously been identified in *S. haematobium* and *S. mansoni* (Campos et al., 2014) and in *S. bovis* and *S. haematobium* hybrids infecting hamster (Mathieu-Bégné et al. 2024). These receptors are responsible for detecting many extracellular signals and transducing them to the G proteins, which then communicate with various downstream effectors such as key molecules involved in developmental and neuromuscular functions (Lagerström & Schiöth, 2008). Transcripts encoding neurotransmitter-sodium-symporter family proteins were also detected. These symporters regulate neurotransmitter uptake, potentially influencing parasite behavior that facilitates the interaction between males and female originating from different species (Rudnick et al. 2014; Mathieu-Bégné et al. 2024).

## Host change related transcriptional plasticity of the parasite *S. bovis*

The change of host can trigger a high remodeling of gene expression in parasites, and there are few works showing that parasite species have shown plasticity at the molecular level that allows them to adapt rapidly to a new host (Hébert et al. 2017; Orsucci et al. 2018; Mathieu-Bégné et al. 2022). We found an intense remodeling of gene expression associated with host change, with 30% of the genes differentially expressed between males *S. bovis* from sheep and hamster, and 26% in females. The sheep represents a completely different habitat for schistosome parasites compared to a hamster, particularly in terms of space, metabolic processes, and blood flow dynamics. These differences can significantly influence the parasites’ life cycle, development, and interaction with the host’s immune system.

Changes in the transcriptomes of males and females show similar patterns in terms of quantity, with 2488 differentially expressed genes (DEGs) in males compared to 2891 in females. Both groups exhibit comparable percentages of over-expressed and under-expressed genes. However, the underlying biological processes differ between the sexes.

In males, we found that over-expressed genes *S. bovis* infecting sheep compared with those infecting hamsters were significantly enriched in action potential, sensory perception, microtubule-based movement, mating, postsynapse organization and reflex. Action potential, sensory perception, postsynapse organization and reflex are part of the same pathway since the nervous system receives an action potential from the sensory neuron cells following an external stimulus triggering a behavior. Neuronal processes have been shown to play a role in male-female worm interactions (Leutner et al. 2013). These biological processes are also critical to colonize and navigate inside the mesenteric system of the definitive host (Nation et al. 2020).

Oxidative phosphorylation, protein targeting, and protein folding were found to be enriched in under-expressed genes in males and females *S. bovis* infecting sheep, these processes are associated to metabolism and energy acquisition (Cardol et al. 2009) and were also found in other parasites using alternative host species suggesting that parasites invest more in these functions to compensate a less efficient use of the host (Mathieu-Bégné et al. 2022). An increase in the expression of genes associated with metabolic pathways may either be a cause— indicating that more energy is required for the parasite to thrive in a larger host—or a consequence—suggesting that resources provided by the host enable the parasites to allocate more energy toward their development. However, paired females are generally retained by the males within their gynecophoric canal, meaning they do not require significant energy to maintain themselves in the blood vessels. Therefore, it is likely that the observed increase in metabolic pathway expression is more of a consequence, with sheep providing a more favorable environment for the development of *S. bovis* females and males.

## Global transcriptomic changes and common biological processes associated with introgression and host change

The number of DEGs between *S. bovis* infecting sheep compared to those infecting hamsters was much higher (30% of the genes) than the number of DEGs we found in introgressed worms (4% of the genes). This suggests that host change induces a higher transcriptomic response in *S. bovis* than an interspecific hybridization event with *S. haematobium*. Host change often requires adaptations at multiple levels, such as coping with altered blood flow, nutrient availability, and host immune responses. These challenges likely drive the extensive transcriptomic reprogramming observed in *S. bovis* when transitioning from a rodent (hamster) to a ruminant (sheep) host, as parasites must rapidly remodel their gene expression to adjust to new biochemical and structural environments. By contrast, the process of introgression, while significant, involves integrating genetic material from closely related species with shared evolutionary histories, such as *S. haematobium* and *S. bovis*. This genetic similarity may limit the need for extensive transcriptomic rewiring, as the hybrids inherit already compatible physiological adaptations to the host environment.

We found 5 common biological processes for which changes in gene expression were found to be associated with the introgression process and the host change. These include the nuclear-transcribed mRNA catabolic process, inner mitochondrial membrane organization, microtubule-based movement, response to endoplasmic reticulum stress and sensory perception. These biological processes have a wide impact on gene function which can allow to cope with environmental changes.

The common enriched biological processes suggest a set of adaptations of *S. bovis* and introgressed worms, that enhance its survival and infectivity across different hosts, contributing to a higher zoonotic potential. Furthermore, the processes observed in hybrid and introgressed worms from sheep are highly similar to those seen in hybrids obtained from hamsters (Mathieu-Bégné et al. 2024). This suggests that there are ’universal’ mechanisms that enable hybridization regardless of the host species. We hypothesized that the enrichment of nuclear-transcribed mRNA catabolic processes enables efficient and flexible gene expression and translation, which can help adapt to other hosts (Wegener and Müller-McNicoll 2018). The energy required to cope with environmental stress is provided by the enrichment of inner mitochondrial membrane organization process (Giacomello et al. 2020). For motility and migration within the host, the enrichment of the movement of microtubules is indispensable (Schroer and Sheetz 1991). The enrichment of the response to endoplasmic reticulum stress may help the parasite evade the host’s immune system (Peng et al. 2021), while the enhancement of sensory perception could facilitate host recognition, avoid immune responses and reach optimal location for mating (Wheeler et al. 2022).

Here we provide insights about gene expression changes and the biological processes involved in introgressive hybridization and host change, both mechanisms allowing schistosome parasites to adapt to new environmental changes facilitating transmission and increasing the zoonotic potential of schistosome hybrids. Further investigation into the role of the biological processes highlighted in this work, and the molecular functions associated, will be essential to understand the full implications of schistosome worm’s transcriptomic plasticity during introgressive hybridization and host change, and how it affects their zoonotic potential.

Understanding the factors contributing to the zoonotic potential of *S. bovis* and its hybrids with *S. haematobium* is crucial for developing more effective prevention and control strategies for urogenital schistosomiasis. Traditionally, intervention efforts have focused on treating infected individuals and controlling snail populations, primarily targeting species that infect humans (Lardans & Dissous, 1998; Touré et al., 2008). However, the presence of introgressed schistosomes, which can display altered transmission dynamics, drug resistance, and host preferences, calls for a reevaluation of these strategies (Leger & Webster, 2017).

Importantly, transcriptomic plasticity in these hybrid parasites allows for colonization of new hosts and environments, facilitating their survival in both human and animal reservoirs. This plasticity not only enhances their zoonotic potential but also complicates control measures by expanding their host range and transmission pathways. Therefore, a deeper understanding of transcriptomic changes in these hybrids is essential for addressing their zoonotic spread and for the development of more targeted intervention strategies (Borlase et al., 2021).

## Data availability

Sequencing data from *S. bovis*, hybrids and introgressed worms infecting sheep will be available at the NCBI-SRA under the BioProject accession number PRJNA984202.

## Acknowledgements

This work has been funded by the French Research National Agency (project HySWARM, grant no. ANR-18-CE35-0001). This study is set within the framework of the “Laboratoire d’Excellence (LABEX)” TULIP (ANR-10-LABX-41). We thank the Bioenvironment sequencing platform of the University of Perpignan.

